# *Achromobacter xylosoxidans* modulates *Pseudomonas aeruginosa* virulence through a multi-target mechanism of competition

**DOI:** 10.1101/2025.02.26.640371

**Authors:** Alison Besse, Quentin Menetrey, Vincent Jean-Pierre, Sylvaine Huc-Brandt, Fabien Aujoulat, Chloé Dupont, Raphaël Chiron, Jean Armengaud, Estelle Jumas-Bilak, Virginie Molle, Lucia Grenga, Hélène Marchandin

## Abstract

The colonization and persistence of *Pseudomonas aeruginosa* in chronically diseased lungs are driven by the production of various virulence factors. However, pulmonary infections in cystic fibrosis (CF) patients are predominantly polymicrobial. While *Achromobacter xylosoxidans* is an opportunistic pathogen in these patients, its impact on *P. aeruginosa* virulence during co-infection remains largely unknown. This study investigated the interaction between *P. aeruginosa* and two clonally related *A. xylosoxidans* strains, Ax 198 and Ax 200, co-isolated from a CF patient sputum. We found that the interaction between *P. aeruginosa* and co-isolated *A. xylosoxidans* was strain-dependent, with the Ax 200 strain significantly reducing *P. aeruginosa* virulence in a zebrafish model, providing the first *in vivo* evidence of interaction between these two species during co-infection. Proteomic analysis revealed that the *P. aeruginosa* proteome was differently impacted by the two *A. xylosoxidans* strains, with Ax 200 altering proteins involved in biofilm formation, swimming motility, iron acquisition, and secretion systems. These proteomic findings were further validated by phenotypic assays, which confirmed that *A. xylosoxidans* affected major *P. aeruginosa* virulence phenotypes, including biofilm formation, swimming motility, and siderophore production. Genetic analysis also confirmed that distinct regulatory mechanisms, including mechanisms involved in the iron cycle, may account for the strain-dependent interaction effects of *A. xylosoxidans* with *P. aeruginosa*. These findings reveal a novel multi-target competitive mechanism through which *A. xylosoxidans* significantly disrupts *P. aeruginosa* virulence.

**IMPORTANCE:** *Pseudomonas aeruginosa*, a major human pathogen, is a leading cause of mortality in cystic fibrosis (CF) patients. Understanding its virulence mechanisms is critical for developing effective infection management strategies. Given the polymicrobial nature of CF infections, it is essential to investigate interspecies interactions that may influence bacterial virulence. While *A. xylosoxidans* is recognized as an opportunistic pathogen in CF, its impact on *P. aeruginosa* virulence during co-infection remains largely unexplored. Deciphering the molecular basis of *P. aeruginosa* virulence in polymicrobial settings could reveal novel and specific therapeutic targets to improve treatment strategies.

## INTRODUCTION

*Pseudomonas aeruginosa* (Pa) is ubiquitous in moist environments like water and soil (1, 2), and is a leading pathogen in healthcare-associated infections (3–6). Pa is also a major cause of lung infection in people with cystic fibrosis (CF) (7). Colonization and persistence of Pa within chronically diseased lungs is driven by the production of virulence factors, such as biofilm formation with increased exopolysaccharide (EPS) production; quorum sensing (QS); motility and attachment involving flagella, type IV pili (T4P) and lectins; secretion systems (TSSs); production of pyocyanin and rhamnolipids; and iron acquisition systems (8). However, pulmonary infections in CF patients are predominantly polymicrobial (9, 10), with the two major pathogens, Pa and *Staphylococcus aureus*, alongside other species such as *Achromobacter xylosoxidans* (Ax), which are increasingly recognized as opportunistic CF pathogens (7, 11). Although co-isolation of Ax and Pa has been reported (12–14), inter-species interaction has rarely been studied, and the impact of Ax on Pa virulence during co-infection remains largely unknown. So far, only two studies have reported *in vitro* interaction between Pa and Ax, including our own previous work (13, 14). We already demonstrated that interactions between Pa and Ax strain pairs significantly inhibited Pa growth, swimming motility, and pigment production (13). In addition, Sandri *et al*. reported cases of either competition or coexistence between strains of these two species (14). Yet, the molecular mechanisms underlying the interaction between them remain unexplored. Furthermore, no *in vivo* studies have addressed how this interaction influences Pa virulence, leaving a critical gap in our understanding of the dynamics of these co-infections.

Based on our previous work, we selected a Pa and two Ax strains, co-isolated from a CF sputum sample. Among these, the Pa II.17 and Ax 200 pair showed the strongest competitive interactions, while the Pa II.17 and Ax 198 pair had a noticeably weaker effect on Pa virulence factors *in vitro* (13). The aim of this study is to i) validate *in vivo* previous *in vitro* observations suggesting that the virulence of Pa is reduced in the presence of Ax, ii) uncover the molecular mechanisms involved in the significant reduction of Pa virulence, and iii) explore why strains of Ax exhibit varying capacities to competitively inhibit Pa virulence.

## RESULTS

### Genomic similarity between *A. xylosoxidans* co-isolated strains Ax 198 and Ax 200

The genomes of Ax 198 and Ax 200 were found to have very similar lengths (6.55 Mb). The two Ax genomes also showed 99.98% and 99.90% of similarity, according to average nucleotide identity based on BLAST and digital DNA-DNA hybridization respectively (Fig. S1); indicating their clonality and supporting the idea that they represent two adaptive variants of the Ax strain colonising the patient.

### *A. xylosoxidans* Ax 200, but not Ax 198, reduces *P. aeruginosa* virulence in a zebrafish infection model

First, we assessed the suitability of the zebrafish bath infection model to evaluate Pa-Ax interactions *in vivo* by examining the survival of zebrafish infected with Pa II.17. A dose-dependent relationship between Pa II.17 and zebrafish mortality was observed, with approximately 20% mortality observed at a low dose 30 hours post-infection (hpi), while 100% mortality occurred at the highest dose within 22 hpi (Fig. 1A), confirming that this is a robust experimental system for studying Pa virulence. On the other hand, both Ax 200 and Ax 198 (suspension at OD_600_=0.5) were avirulent, causing no mortality after 30 hours of bath immersion, similar to the control group (Fig. 1B). The impact of Ax 200 and Ax 198 on Pa virulence was then evaluated. Pre-immersion with Ax 200 (OD_600_=0.5) prior to introducing Pa II.17 (OD_600_=0.2), significantly reduced embryo mortality, with 50% surviving beyond 30 hpi, compared to 100% mortality at 22 hpi when infected with Pa II.17 alone (Fig. 1B). In contrast, pre-immersion with Ax 198 (OD_600_=0.5) resulted in mortality levels comparable to those observed with Pa II.17 alone (Fig. 1B).

**Figure 1.**
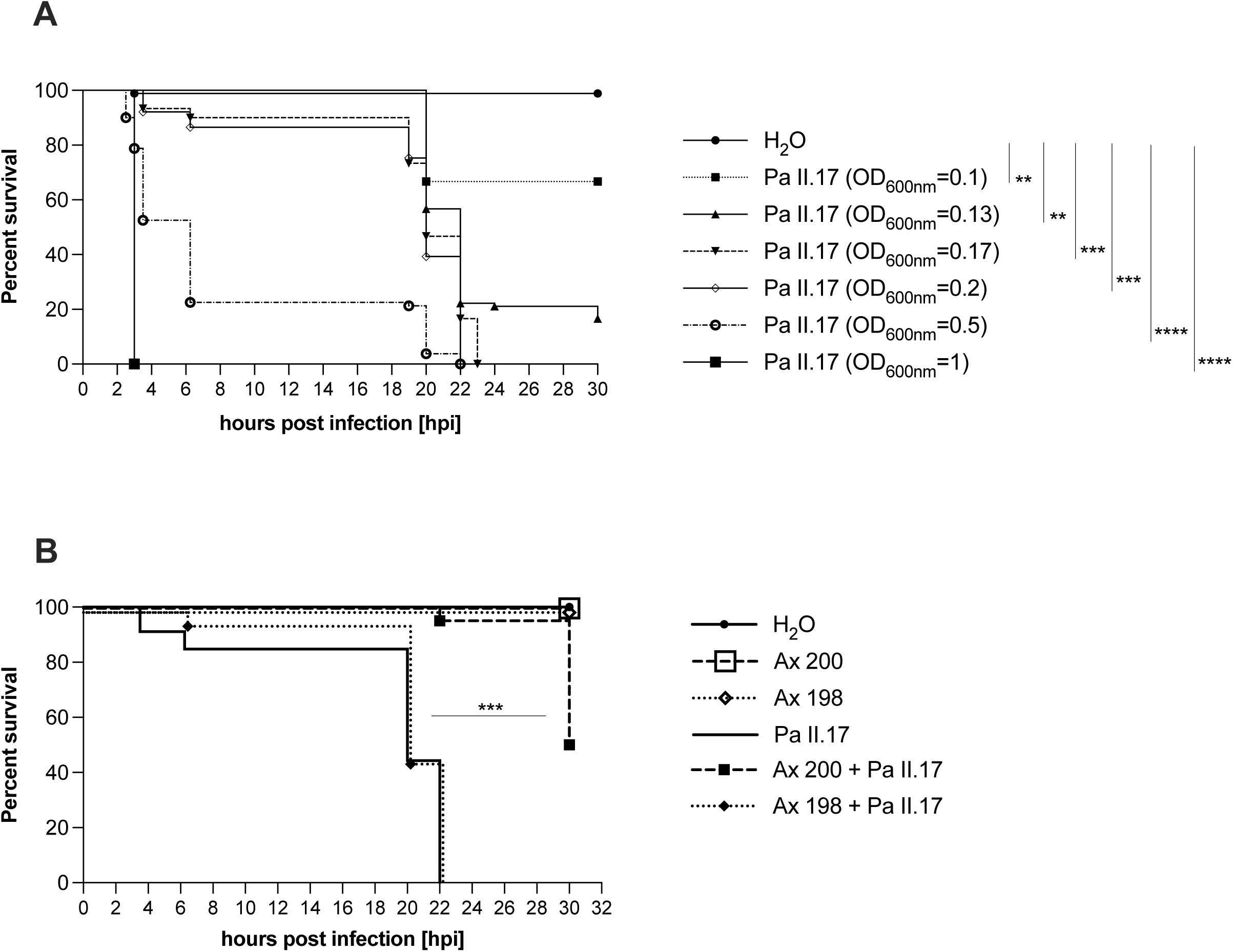
Survival outcomes in the zebrafish infection model with Pa II.17 alone (A) or co-infections with Ax 200 or Ax 198 (B). Survival of zebrafish embryos at 48 h post-fertilization is represented as a Kaplan– Meier plot. (A) Zebrafish embryos were infected by bacterial suspensions of Pa II.17 at different concentrations (OD_600_ of 0.1, 0.13, 0.17, 0.2, 0.5 and 1, corresponding to concentrations ranging from 3x10^8^ to 4x10^9^ CFU/mL); (B) Zebrafish embryos were infected by bacterial suspensions of either Pa II.17 (prepared in fish water at an OD_600_ of 0.2 corresponding to approximately 6x10^8^ CFU/mL), Ax 200 (prepared in fish water at an OD_600_ of 0.5 corresponding to approximately 1x 10^9^ CFU/mL) or Ax 198 (prepared in fish water at an OD_600_ of 0.5 corresponding to approximately 1x 10^9^ CFU/mL) strains alone, or with Ax 200 or Ax 198 preincubation of 1.5 h before Pa II.17 introduction. Embryos maintained in fish water were used as negative control. Results are presented as the proportion of surviving embryos (n > 20 for each, indicative of at least two separate experiments). Significant differences tested by log-rank test are indicated (**, *P* ≤ 0.01; ***, *P* ≤ 0.001; ****, *P* ≤ 0.0001).

The proteome of *P. aeruginosa* Pa II.17 is more significantly impacted in co-culture with *A. xylosoxidans* Ax 200 than with Ax 198

When co-cultured with Ax 200, 623 Pa proteins out of the 2019 detected showed significant differential abundance compared to the Pa II.17 monoculture (p-value ≤ 0.05 and log2FC |1|, Fig. 2A, Fig. S2A and S2B, Tables S1 and S2). All of these proteins exhibited a log2 FC of less than -1, indicating a significant reduction in abundance during co-culture (Fig. 2A). In contrast, co-culture with Ax 198 resulted in only 35 out of 2216 detected proteins showing significant differential abundance (Fig. 2B, Fig. S2A and S2B). Among these, only 13 proteins were similarly less abundant in both Ax 198-Pa II.17 and Ax 200-Pa II.17 co-cultures, suggesting distinct molecular mechanisms underlying the interactions between Pa II.17 and the two Ax strains. Proteins with significant changes in abundance in co-cultures were predominantly classified into categories “Translation, ribosomal structure and biogenesis”, “Energy production and conversion” and “Amino acid transport and metabolism”. Several were also grouped into the “Unknown function” category (Fig. S2B). Notably, the latter included key proteins, such as PelC and AlgZ, involved in EPS biosynthesis, and the outer membrane porin OprD, known for its roles in the transport of amino acids and the uptake of antibiotics (15).

**Figure 2.**
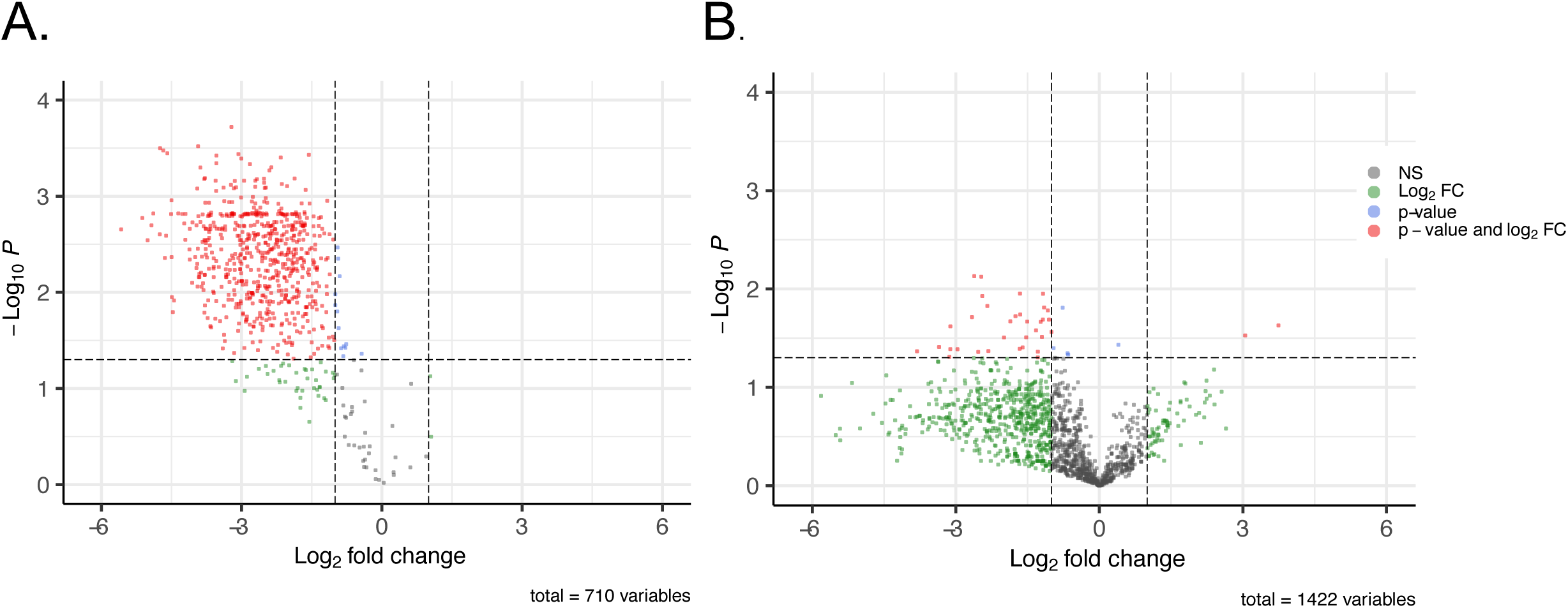
*P. aeruginosa* Pa II.17 proteins displaying differential abundance between mono-culture and co-culture with *A. xylosoxidans* Ax 200 (A) or *A. xylosoxidans* Ax 198 (B). Each dot in the Volcano Plot represents a protein identified both in mono- and co-culture. Each mono- and co-cultures was analysed in triplicate. Changes in protein abundance were considered statistically significant (red dots) when the P-value was ≤0.05 (Student’s t-test, horizontal dashed line) and the log2 FC exceeded |1|. NS = not significant.

Some Pa II.17 proteins were also exclusively detected either in mono- or co-culture (Fig. S2C), the majority of them (99.4% and 72%) being exclusively detected in the Pa monoculture and absent in the Ax 200-Pa II.17, or the Ax 198-Pa II.17 co-culture, respectively (Tables S1 and S2).

By combining the proteins with differential abundance and those exclusively detected in either mono- or co-culture, the total number of Pa proteins impacted was 1936 for the co-culture with Ax 200 and 831 for the co-culture with Ax 198. Notably, only 410 of these 831 proteins (49.3%) overlapped with those affected in the Ax 200-Pa II.17 co-culture, further highlighting distinct interaction mechanisms (Tables S1 and S2).

Altogether, Ax 200 exerts a more pronounced negative impact on Pa II.17 proteome, potentially affecting Pa growth, proliferation, metabolism, and most likely, virulence to a greater extent than Ax 198; and molecular interactions between Pa II.17 and Ax 200 are mechanistically distinct from those with Ax 198. The following sections focus on the main Pa virulence factors inhibited by Ax.

Biofilm formation and swimming motility in *P. aeruginosa* Pa II.17 is severely impaired in co-culture with *A. xylosoxidans* Ax 200

Key Pa virulence proteins were underrepresented or absent in the presence of Ax 200. Specifically, T4P proteins, including PilG, PilU, PilM, FimL, and FimV, were at least three times less abundant in Ax 200-Pa II.17 co-culture compared to Pa monoculture (Fig. 3, Table S1). As T4P is essential for twitching motility (16–19), this suggests that Pa adhesion may be impaired during co-culture. In addition, PelC, involved in EPS production, was four times less abundant in co-culture (Fig. 3). As EPS secretion contributes to biofilm structural integrity, later stages of biofilm formation could also be negatively impacted by Ax 200. Moreover, FleQ, the master regulator of biofilm formation, was nearly 14 times less abundant in co-culture, further supporting that biofilm formation could be defective. Also, other proteins associated with biofilm regulation, such as SiaB, or with T4P formation such as PilT, PilB and PilD, were exclusively detected in Pa mono-culture (Fig. 3). To validate whether biofilm formation was indeed impaired by Ax 200, a biofilm formation assay was conducted. Consistent with our proteomic analysis, *in vitro* biofilm biomass was two times lower in Ax 200-Pa II.17 co-culture compared to Pa monoculture (Fig. 4A).

**Figure 3.**
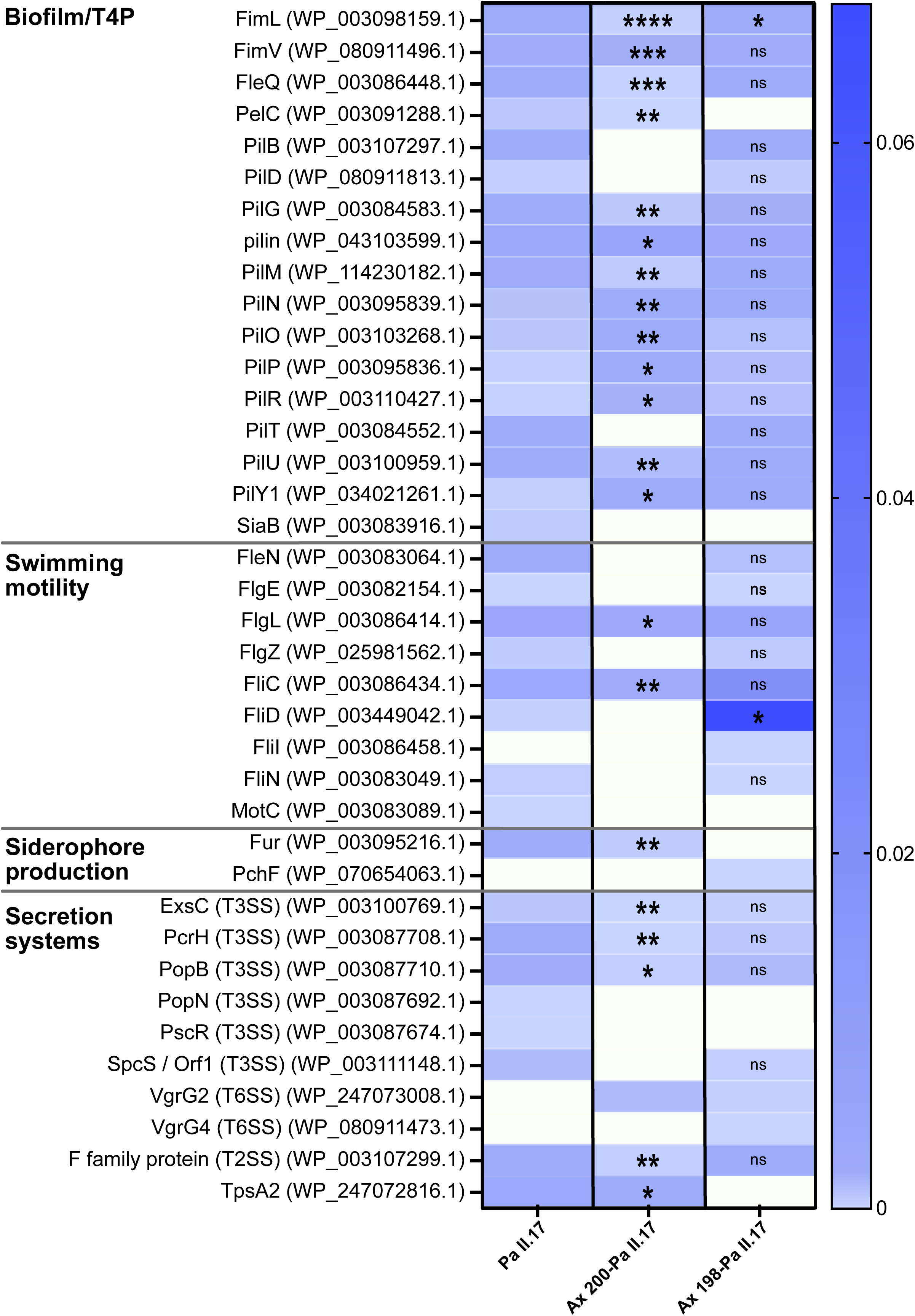
Abundance of *P. aeruginosa* Pa II.17 virulence-associated proteins in monoculture and co-culture with *A. xylosoxidans* Ax 200 or Ax 198. Protein annotations were verified using BLASTp. Shades of blue indicate the average Normalized Spectral Abundance Factor of three replicates. Only proteins with a significant difference in abundance between mono- and co-culture and proteins exclusively detected either in mono- or co-culture are shown. The statistical significance of abundance variation between Pa II.17 mono-culture and Ax 200-Pa II.17 or Ax 200-Pa II.17 co-culture was assessed using a Student t-test; ****, *P* ≤ 0.0001; ***, *P* ≤ 0.001; **, *P* ≤ 0.01; *, *P* ≤ 0.05; ns = not significant. Blank cells indicate proteins not detected for which no statistical test was performed.

**Figure 4.**
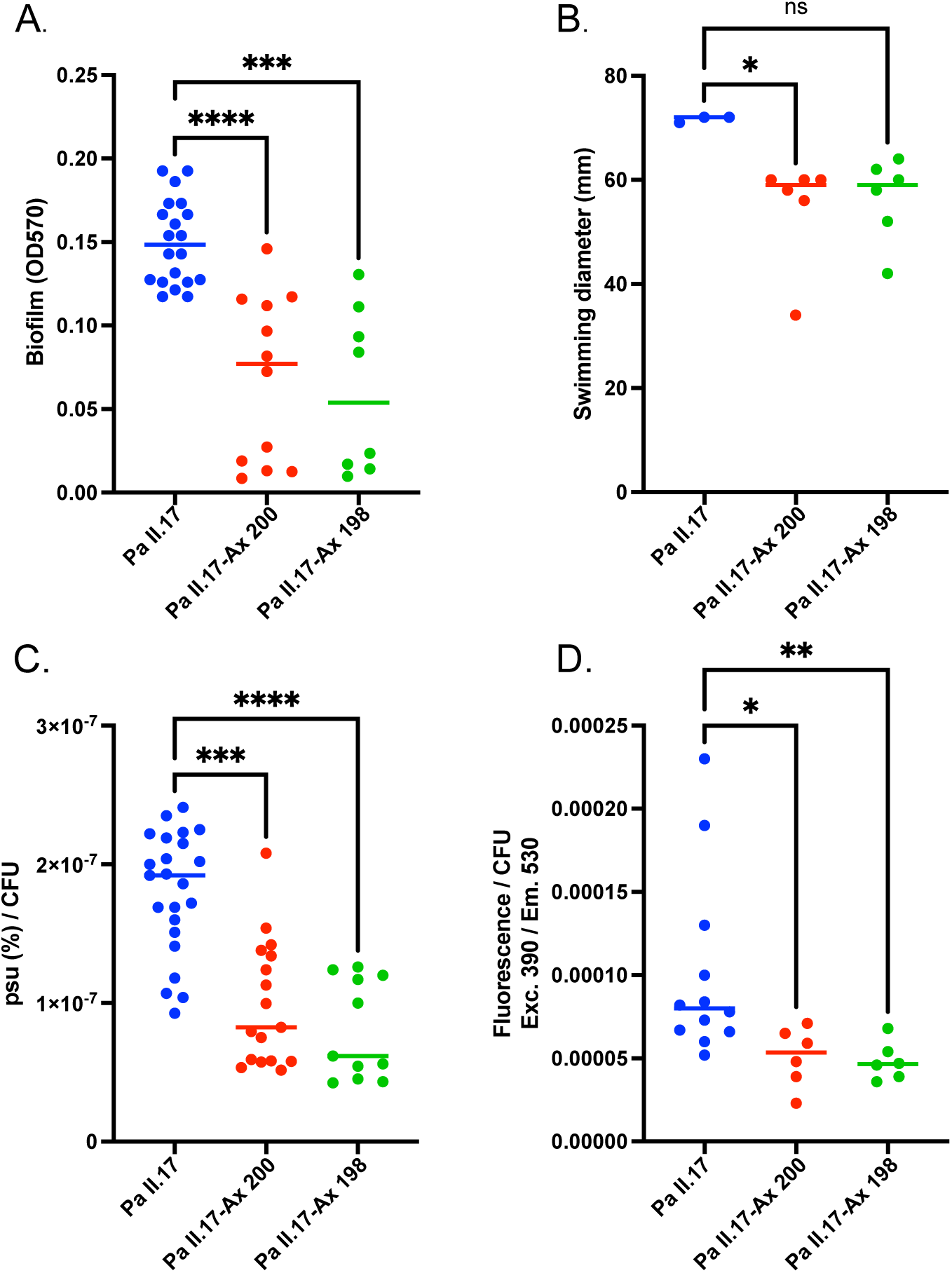
Biofilm formation (A), swimming motility (B), siderophore production (C) and pyoverdine production (D) of *P. aeruginosa* Pa II.17 in monoculture and co-culture with *A. xylosoxidans* Ax 200 or Ax 198. The values are medians. Phenotypic tests were normalised between assays. Siderophore production was normalised on total CFU since *A. xylosoxidans* and *P. aeruginosa* are both able to produce siderophores. Pyoverdine production was normalized exclusively on *P. aeruginosa* CFU. The statistical significance of the results was calculated by a nonparametric Kruskal-Wallis test, ****, *P* ≤ 0.0001; ***, *P* ≤ 0.001; **, *P* ≤ 0.01; *, *P* ≤ 0.05; ns = not significant.

Pa II.17 swimming motility which is crucial for both the initial adhesion phase and the dissemination of biofilms (20) is likely to be also impacted by Ax 200, as flagellar proteins were found to be 1.7 to 2.2 times less abundant in co-culture (Fig. 3). These include FlgL, which is inserted in the hook-filament junction; and the filament flagellin FliC. Several other proteins related to flagellum formation or function were detected only in Pa II.17 monoculture (Fig. 3). A motility assay confirmed that swimming was significantly reduced by 1.22-fold in co-culture compared to monoculture, consistent with the reduced abundance of flagellar proteins (Fig. 4B).

### Iron uptake by P. aeruginosa is impaired in co-culture with *A. xylosoxidans* Ax 200

The ferric uptake regulator protein Fur, which is essential for maintaining iron homeostasis, was seven times less abundant in Ax 200-Pa II.17 co-culture compared to Pa monoculture (Fig. 3, Table S1). Additionally, the pyochelin receptor FptA and another siderophore receptor were exclusively detected in Pa monoculture (Table S1). These proteins display a signal peptide for secretion and are thus potentially secreted (21). Iron uptake in Pa II.17 might thus be negatively affected by the presence of Ax 200, prompting further investigation in optimal conditions for assessing siderophore production impairments, *i.e*., using the iron-depleted medium MM9 (22). In this condition, total siderophore production was reduced by nearly half in the co-culture (Fig. 4C) and pyoverdine production, one of the primary Pa siderophores, was also drastically reduced (Fig. 4D).

### Abundance of secretion system proteins is reduced in *P. aeruginosa* co-cultured with *A. xylosoxidans* Ax 200

Proteins associated with the Type III secretion system (T3SS), such as PopB, a translocator protein essential for pore formation; PcrH, a chaperone stabilizing effector protein; and ExsC, a scaffold protein involved in T3SS assembly, were less abundant in co-culture with Ax 200. Additionally, numerous other structural and regulatory proteins of T3SS were exclusively detected in monoculture (Fig. 3, Table S1). A protein associated with the Type II secretion system (T2SS) was also less abundant in co-culture, suggesting a reduced efficiency of T2SS in the presence of Ax 200. Similarly, the Type VI secretion system (T6SS) was significantly impacted, with several proteins detected only in monoculture, indicating impaired assembly or function of T6SS in co-culture (Fig. 3, Table S1). Interestingly, VgrG2 was the only T6SS-protein found exclusively in co-culture and not in monoculture. However, the underrepresentation of other essential T6SS components in co-culture suggests a compromised functionality of T6SS in the presence of Ax 200.

### Proteomic changes are limited in *P. aeruginosa* co-cultured with Ax 198, but key virulence phenotypes are still affected

In contrast to Ax 200, the presence of Ax 198 in co-culture with Pa II.17 did not significantly reduce the abundance of proteins involved in biofilm, T4P or TSS system functionality (Fig. 3). Indeed, only FimL, a central regulator required for T4P biogenesis, biofilm development, and T3SS function, was less abundant in Ax 198-Pa II.17 co-culture compared to Pa monoculture. This suggested that specific Pa phenotypes could still be affected in the presence of Ax 198. Phenotypic assays revealed that biofilm formation by Pa II.17 was significantly reduced in co-culture with Ax 198 (Fig. 4A). This reduction might also be explained by the absence or reduction of several proteins exclusively detected in Pa monoculture, including SiaB and PelC for biofilm regulation; PopN and PscR for T3SS; and LcmF2, ClpV2, Hcp and HsiC2 for T6SS (Fig. 3, Table S2).

Regarding the abundance of QS proteins, none of the changes detected between Pa monoculture and Ax 198-Pa II.17 co-culture were statistically significant in our conditions. However, some QS proteins such as the acyl-homoserine lactone (AHL) synthetase LasI, and third QS system proteins PqsB and PqsE, were not detected in Ax 198-Pa II.17 co-culture (Table S2). Overall, QS proteins exhibited greater alterations in the Ax 198-Pa II.17 co-culture compared to Ax 200, where no differences were observed between mono- and co-culture.

Regarding swimming motility, only FliD, a flagellar cap protein, was more abundant in Ax 198-Pa II.17 compared to monoculture (Fig. 3). This overabundance alone is unlikely to enhance swimming motility. In addition, three flagellar structural proteins, FlgG, FlaG and FliI, were detected exclusively in Ax 198-Pa II.17 co-culture (Table S2), suggesting potentially more efficient flagellar formation. Nonetheless, phenotypic assays showed no significant differences in swimming motility for Ax 198-Pa II.17 co-culture compared to monoculture (Fig. 4B). This could be explained by the absence of essential regulators or structural proteins, such as FliH, FliS, MotC and FlgN, which were not detected in co-culture (Table S2).

Concerning iron acquisition, siderophore production in Ax 198-Pa II.17 co-culture was significantly reduced compared to monoculture, similar to the reduction observed with Ax 200 (Fig. 4C). Notably, proteomic data highlighted distinct profiles between the two conditions. For example, while Fur was less abundant in co-cultures with Ax 198 or Ax 200, PchF, required for pyochelin biosynthesis, was only detected in Ax 198-Pa II.17 co-culture (Fig. 3).

### Genetic basis for differential effects of *A. xylosoxidans* Ax 198 and Ax 200 on *P. aeruginosa* virulence

Sixty variations, either single nucleotide polymorphisms (SNPs) or deletions/insertions, were identified between the Ax 200 and Ax 198 genomes, of which 37 were located in coding sequences corresponding to 31 genes (Table S3). Most of these genes encode proteins of “unknown function”, while the remaining genes encode proteins primarily involved in transcription (Fig. S3). Based on Bakta and Prokka annotations, SNPs were identified in three genes encoding transcriptional regulators: *dmlR*, *nusA* and a tetR-type helix-turn-helix (HTH) domain-containing protein-encoding gene (Table S3). The corresponding transcriptional regulators regulate pyruvate metabolism, transcript elongation, and tetracycline resistance, respectively. Additionally, five SNPs were identified in the *yqjI* gene of Ax 200, leading to the loss of a start codon and subsequent absence of functional YqjI protein production (Table S3). YqjI, a PadR family transcriptional regulator, is known to control the transcription of *yqjH* in *Escherichia coli* (23), which encodes the ferric reductase YqjH, a key enzyme promoting reduction of Fe^3+^ and its release from siderophores in the cytoplasm. Furthermore, a SNP was found in the ferripyoverdine receptor FhuE-encoding gene between the Ax 198 and Ax 200 genomes. Also, in the Ax 200 genome, a SNP caused the loss of a stop codon in a porin-encoding gene, suggesting a potentially dysfunctional porin. Notably, no SNPs were detected in genes encoding TSS components or regulators. These findings suggest that distinct regulatory mechanisms, including those related to iron metabolism, contribute to the differences in Ax-Pa competition between Ax 198 and Ax 200.

## DISCUSSION

As lung infections in CF patients are considered polymicrobial (24), investigating interbacterial competition and its effects on virulence is particularly relevant. Although the major CF pathogen Pa is increasingly exposed to Ax during polymicrobial lung infection in CF patients (25), the most frequently isolated species (26, 27), the interaction between these two species remains poorly understood.

### Comprehensive mechanisms of *P. aeruginosa*-*A. xylosoxidans* competition: Ax 200 disrupts multiple Pa virulence pathways, including biofilm formation, secretion systems, and iron uptake

The colonization and persistence of Pa in the respiratory tract rely on its remarkable versatility including transitioning from a planktonic motile lifestyle during host invasion to biofilm formation, enabling immune evasion and antimicrobial resistance (28). This transition is tightly regulated by the intracellular second messenger c-di-GMP and its receptor-effector, FleQ, a master transcriptional regulator. In response to high c-di-GMP, FleQ represses flagellar biosynthesis and activates EPS production genes (*psl*, *pel*, *cdr*AB), enhancing biofilm formation by inhibiting swimming motility and promoting matrix production (29–31). FleQ detected in Pa II.17 proteome was significantly reduced in co-culture with Ax 200, likely altering the transcription of FleQ-dependent genes (30). Moreover, an essential feature of the initial stage of Pa biofilm development is surface adhesion *via* twitching motility (32), mediated by T4P extension and retraction (33). PilG and PilU, key T4P proteins (16, 34), were significantly less abundant in the Pa II.17 proteome during co-culture with Ax 200 likely contributing to the observed reduction in biofilm formation. By uncovering the molecular basis for the reduction of biofilm in co-culture, we demonstrated that Ax 200 not only impairs biofilm formation (from initial adhesion and matrix production to dissemination *via* swimming motility) but also disrupts the planktonic-to-sessile transition, thereby limiting Pa adaptive capacity. T4P is also a critical adhesin facilitating host epithelial cell colonization (18). The reduced abundance of T4P structural and functional proteins in Ax 200-Pa II.17 co-culture likely contributes to diminished host cell infection, as evidenced by the lower zebrafish mortality observed during co-infection. Therefore, our study validates and extends the findings of two previous studies that reported reductions in biofilm formation and motility *in vitro* for this strain pair and other Ax-Pa combinations (13, 14). By incorporating *in vivo* experiments and proteomic analyses, our work goes beyond these earlier studies, providing deeper insights into the molecular mechanisms underlying these phenotypic changes and demonstrating their relevance in a host infection model.

Secretion systems are key virulence factors of Pa (18, 35). FimL is a central regulator required not only for T4P activity and biofilm development, but also for the functionality of T3SS that enables Pa to inject effectors into host cells, disrupting their machinery, inducing cytotoxicity, and enhancing bacterial survival (36). Our findings showed that FimL was less abundant in Pa when co-cultured with Ax 200, and as expected, the T3SS-associated proteins were also less represented. This is the first report of T3SS disruptions during Pa-Ax interactions*. P. aeruginosa* possesses four other TSSs (35). T2SS was also affected in Pa-Ax co-culture with several T2SS-related proteins being less abundant in co-culture. T2SS+ Pa strains are known to cause lethal infections in mice, albeit more slowly than T3SS+ strains (37). Finally, T6SS, known to mediate Pa internalization into eukaryotic cells via effectors such as Vgr2 (38), was significantly impaired, indicating that Ax 200 disrupts T6SS functionality in Pa during co-culture.

The Pa virulence arsenal also includes siderophores secreted to facilitate iron uptake (39), which is essential for numerous cellular processes. During infection, Pa competes with the host and microbial species for iron, relying on two siderophores, pyoverdine and pyochelin, which are critical for establishing successful infections (8, 40). For example, siderophores are central to the interplay between the two main CF pathogens, Pa and *S. aureus*, as they are required for Pa to kill *S. aureus* efficiently (41, 42). However, no prior studies have reported the involvement of iron homeostasis or siderophore production in interactions between Pa and *Achromobacter.* Here, we observed that siderophore production by Pa II.17 was significantly reduced when co-cultured with Ax 200. Since Ax is also capable of producing siderophores (43), it was initially unclear which microorganism was responsible for this decrease. However, by measuring pyoverdine, we confirmed that Pa was at least partially responsible for the observed siderophore reduction. Moreover, Pa has more than 30 receptors that recognize its own siderophores as well as xenosiderophores produced by other bacteria, facilitating their transport into the cells (44). Our proteomic analyses revealed that at least two receptors, including the pyochelin receptor FptA, were detected in Pa monoculture only, suggesting that under our experimental conditions where iron was not limited, Pa iron uptake might be less effective in the presence of Ax 200. Similarly, the Fur regulator, critical for iron homeostasis, was significantly less abundant in the presence of Ax 200 (45, 46). This is the first study to report the involvement of iron homeostasis and siderophore production in Ax-Pa interactions, suggesting that this may be a key factor in attenuating Pa virulence. Supporting this hypothesis, we found that the ferrireductase regulator-encoding gene *yqjI* was mutated in the Ax 200 genome.

Altogether, this study highlights a multi-target mechanism of competition between Ax and Pa, whereby Ax simultaneously disrupts multiple Pa virulence factors. This finding challenges the prevailing view of Pa as a highly competitive species that outcompete other species (47, 48). Moreover, the patient colonization history, marked by chronic Ax colonization and sporadic Pa presence, raises questions about the role of bacterial competition in shaping infection outcomes (49). Examining the effect of Pa on the Ax proteome could provide further insights into these dynamics.

### Strain variability in *A. xyloxidans* determines its effect on *P. aeruginosa* virulence

Our findings reveal that interaction between Pa and Ax is strain-dependent, likely driven by distinct mechanisms specific to each strain. A previous study on strains isolated from the same patient at different infection stages has also described varied interactions between Pa and Ax, ranging from coexistence and competition (14). Our study is the first to report distinct types of interaction between Pa and Ax strains co-isolated from the same sputum sample highlighting how adaptive microevolution in CF lungs can generate sub-clonal co-existing Ax variants with differing competitive abilities against Pa. Such diversity within the Ax population likely provides an adaptive advantage, enabling the population to respond to environmental changes, including new colonization episodes by opportunistic pathogens like Pa, and supporting the long-term survival of Ax in CF lungs (49–52). Regarding underlying mechanisms, the most notable genomic variations between Ax 198 and Ax 200 were related to transcriptional regulators, iron uptake, and porin genes. In addition, QS regulation may play a role in the interaction with Ax 198 but not with Ax 200. Our findings also excluded that defective TSSs in Ax 198 explain its lesser impact on Pa virulence. Indeed, both Ax 198 and Ax 200 carry genes encoding T6SS, T2SS and T3SS, as well as components of T1SS and T4SS, as reported in CF Ax strains (53–55) and no significant differences in TSS-related genomic regions, including those related to transcriptional, post-transcriptional, or post-translational regulators, were found between both genomes (56). This contrasts with previous studies demonstrating that Ax T6SS could target and kill the Pa reference strain PAO1 (57) and that T6SS VgrG and T1SS components were encoded by a competitive *Achromobacter* strain and absent in a co-existing strain (14).

## MATERIALS AND METHODS

### Patient, bacterial strains and ethical statement

Pa II.17, Ax 198 and Ax 200 were co-isolated from a sputum sample obtain from a CF patient attending the CF center at Montpellier University Hospital, France. The patient was chronically colonised by Ax and sporadically colonized by Pa. Ax 198 and Ax 200 are clonally-related according to previous genotyping, including *nrdA* gene sequencing, Multi-Locus Sequence Typing, Pulsed-Field Gel Electrophoresis and Multiplex rep-PCR (13). This study was approved by the Institutional Review Board at Nîmes University Hospital (IRB no. 19.02.01).

### Bath immersion-based zebrafish embryo survival assay and ethical statement

Overnight bacterial cultures grown at 37°C in Trypticase Soy Broth (TSB) were centrifuged at 3500g for 10 minutes and resuspended in fish water (distilled water with 60 µg/mL sea salt, Instant Ocean, and 4.10^-4^ N NaOH). Bacterial suspensions were adjusted to an optical density of 600 nm (OD_600_), with bacterial counts verified by plating on Tryptic Soy Agar (TSA). Experiments were conducted in fish water at 28°C using the zebrafish model (*Danio rerio*) (AB zebrafish line). Embryos were dechorionated at 48 h post-fertilization (hpf) and infected via bath immersion. Groups of 10 healthy embryos were placed in 6-well plates containing the bacterial suspension and dead embryos were visually identified by the absence of a heartbeat. All experiments were performed in accordance with European Union guidelines for the care and use of laboratory animals (http://ec.europa.eu/environment/chemicals/labanimals/homeen.htm) and were approved by the Direction Sanitaire et Vétérinaire de l’Hérault and the Comité d’Ethique pour l’Expérimentation Animale (CEEA-LR-13007). At the end of experiments, plates were sealed with parafilm, frozen at −20°C for 48 h to ensure the embryo’s death, and autoclaved.

### Mono- and co-culture conditions

Overnight cultures in TSB were used to inoculate mono-cultures (Ax 198, Ax 200, or Pa II.17) or co-cultures (Ax 198-Pa II.17 or Ax 200-Pa II.17) in specific media, depending on the experiment. To account for Ax’s slower growth rate compared to Pa, co-culture conditions followed the protocol of Menetrey *et al*. (13). Briefly, Ax was inoculated firstat OD_600_=0.005, followed by Pa inoculation at OD_600_=0.001 after a 4 h delay. After 48 h of incubation at 37°C, the cultures were used for proteomic analysis or phenotypic assays. In parallel, bacterial cell counts (CFU/mL) were determined using TSA plates and the EasySpiral Pro (Interscience®) system, following the manufacturer’s instructions. Pa and Ax colonies were visually distinguished.

### Proteomic analysis

For each condition (Pa II.17-Ax198 or Pa II.17-Ax 200 co-culture, Pa II.17 mono-culture in two independent assays), proteome extraction was performed as described previously (58). Cultures were centrifugated at 3000g for 15 min at 20°C; pellets were resuspended in 1 ml of PBS 1X pH 7.0 (Sigma-Aldrich), centrifuged at 10000g for 3 min, and subsequently dissolved in 100 μL LDS 1X supplemented with 5% β-mercaptoethanol. Secreted proteins were analyzed after trichloroacetic acid (TCA) (Sigma-Aldrich) precipitation of the culture supernatants, and since most exoproteins were recovered in proteome due to cell lysis, the results from both fractions were combined for analysis. Peptides were analysed using an ESI-Q Exactive HF mass spectrometer (ThermoFisher Scientific) coupled with an Ultimate 3000 Nano LC System (ThermoFisher Scientific). Two μl of peptides were injected onto a reverse phase Acclaim PepMap 100 C18 column (3 μm, 100 Å, 75 μm id × 500 mm) and resolved at 0.2 μL/min with a 90-min gradient of CH_3_CN (4% to 40%) containing 0.1% HCOOH. The tandem mass spectrometer operated in data-dependent mode using a top-20 strategy, selecting peptide molecular ions with double or triple positive charges for fragmentation, with a 10s dynamic (59).

The tandem mass spectrometry (MS/MS) spectra were interpreted using MASCOT Daemon 2.6.0 software (Matrix Science) with genomes of Ax 200, Ax 198, and *Pa* II.17 (accession numbers GCA_022976495.1, GCA_022976515.1, and GCA_022976545.1, respectively). Parameters included full-trypsin specificity, a maximum of one missed cleavage, 5 ppm mass tolerance on parent ions, and 0.02 Da on MS/MS. Modifications considered were carbamidomethylated cysteine as a static modification and oxidized methionine as a dynamic modification. Peptides with MASCOT scores below a *P*-value of 0.05 were included.

Proteins quantification was based on spectral counts using the Normalized Spectral Abundance Factor (NSAF) (60). Each condition included three biological replicates. Proteins were considered detected if MS/MS-assigned spectra were counted in at least two of these replicates. Fold change was calculated as the NSAF ratio of co-cultures to the summed NSAF of Pa mono-cultures. Statistical significance of abundance variation between mono- and co-culture was assessed using a Student t-test. Proteins with statistically non-significant results, or inconsistent abundances between two independent Pa mono-cultures were excluded. Functional annotation of detected proteins was conducted using the eggNOG v5 database (61).

### Quantification of siderophore and pyoverdine production

Mono- and co-cultures were performed in minimal medium MM9 (0.3g/L KH_2_PO_4_, 0.5g/L NaCl, 0.1g/L NH_4_Cl supplemented with 3.3% Casamino acid, 0.2% glucose, 1mM MgCl_2_, 100µM CaCl_2_ and 7.5 mg/L tryptophan) under static conditions according to Payne’s method with modifications (62). Siderophore production was assessed using the universal Chrome Azurol Sulphonate (CAS) assay (62, 63). Briefly, after 48h at 37°C, 80 µL of supernatants were mixed with 80 µL of CAS reagent. Absorbance at 630 nm was measured after 5 min. Siderophore production was expressed as percent siderophore units (psu) calculated using the formula: [(Ar - As)/Ar]100 = % siderophore units, where Ar represents the absorbance of the reference (CAS solution and uninoculated broth) and As represents the absorbance of the sample (CAS solution with the supernatant). To quantify pyoverdine production, 100 μL of culture supernatants were transferred to black 96-well plates wells (Greiner) and fluorescence was measured at excitation/emission wavelengths of 390 nm/530 nm using a multimode microplate reader (TECAN, spark) (64).

Results were normalized based on CFU for each sample. Each assay was performed in triplicate and repeated at least twice. Following confirmation of data normality with the Shapiro-Wilk test, statistical significance was determined using nonparametric Kruskal-Wallis test.

### Quantification of biofilm formation

Biofilm formation was assessed as described previously (13). For dual-species cultures (Ax 198-Pa II.17 and Pa II.17-Ax 200), suspensions prepared in TSB from O/N cultures (initial OD_600_=0.005 for Ax strains and 0.001 for Pa), were mixed in a 1:1 ratio. Biofilm quantification was performed after 48 h of incubation at 37°C based on Harvey *et al*. (65), with modifications (13). The wells were washed thrice with tap water and stained with 1% crystal violet (CV) solution. The CV was then solubilized using 200 μL of 95% ethanol, and 125 μL of the solution was transferred to a new plate for absorbance measurement at 570 nm. Data normality was verified using the Shapiro-Wilk test, and statistical analyses were conducted using one-way analysis of variance (ANOVA).

### Swimming motility assays

Swimming motility was assessed following a modified protocol from Menetrey *et al.* (13). A 2.5 µL aliquot of Ax 198 or Ax 200 suspension at OD_600_=0.5 was inoculated into the swim plates (20 g/L Luria Bertani broth, 0.3% agar). After 4 h of incubation at 30°C, a 2.5 µL aliquot of Pa II.17 suspension at OD_600_=0.1 was inoculated 1.5 cm away from the Ax spot. Plates were incubated at 30°C for an additional 44 h. Pa swimming ability was evaluated by measuring the diameter of the turbid circular zone and comparing it to the zone formed by Pa when cultured alone. All experiments were performed in triplicate.

### Whole genome sequencing and comparison of *Achromobacter* genomes

Bacterial DNA was extracted using the MasterPure extraction kit (Epicentre) and sequenced on an Illumina NextSeq 500 at the Plateforme de Microbiologie Mutualisée (P2M, Institut Pasteur, Paris, France). Reads were assembled *de novo* using SPAdes v3.12.0 (66); contigs were annotated via the NCBI Prokaryotic Genome Annotation Pipeline (PGAP) (67). Ax genome alignments were visualized using BRIG. Sequence Types (ST) were determined via the PubMLST database (https://pubmlst.org). Both substitutions and insertions/deletions (indels) between *Achromobacter* genomes were identified using Snippy v4.6.0 (https://github.com/tseemann/snippy), with core genome SNPs (Single Nucleotide Polymorphisms) determined using Snippy-core. These tools were run on the Galaxy platform (68) with default settings. Functional annotation of SNPs-associated genes was conducted using the eggNOG v5 public database (61). Average Nucleotide Identity by BLAST was calculated using the JSpecies webserver (http://jspecies.ribohost.com/jspeciesws) (69), while digital DNA–DNA hybridization was estimated with GGDC 3.1 (70).

### Data availability

Mass spectrometry proteomic data have been deposited in the ProteomeXchange Consortium via the PRIDE (71) partner repository with the dataset identifiers PXD058911 and 10.6019/PXD058911. Raw genome sequencing data and assemblies were deposited in GenBank under the BioProject accession numbers PRJNA823997 and PRJNA824006.

## ACKNOWLEDGEMENTS

This research received no specific grant from any funding agency in the public, commercial, or not-for-profit sectors. The authors thank Teresa Sawyers, Medical Writer at the B.E.S.P.I.M., Nîmes University Hospital, for her assistance in editing this manuscript, and Corentin Escobar and Caroline Santer, for their contribution to preliminary zebrafish experiments and phenotypic assays.

## AUTHORS CONTRIBUTION

Conceptualization: QM, VM, LG, HM. Data curation, Formal analysis, Investigation, Methodology, Validation: AB, QM, VJP, SHB, FA, CD, LG. Project administration, Supervision: HM. Resources: RC, EJB, JA, VM. Vizualisation: AB, QM. Writing: original draft: AB, QM, HM. Writing: review & editing: all authors.

## SUPPLEMENTARY FIGURES

**Figure S1.**
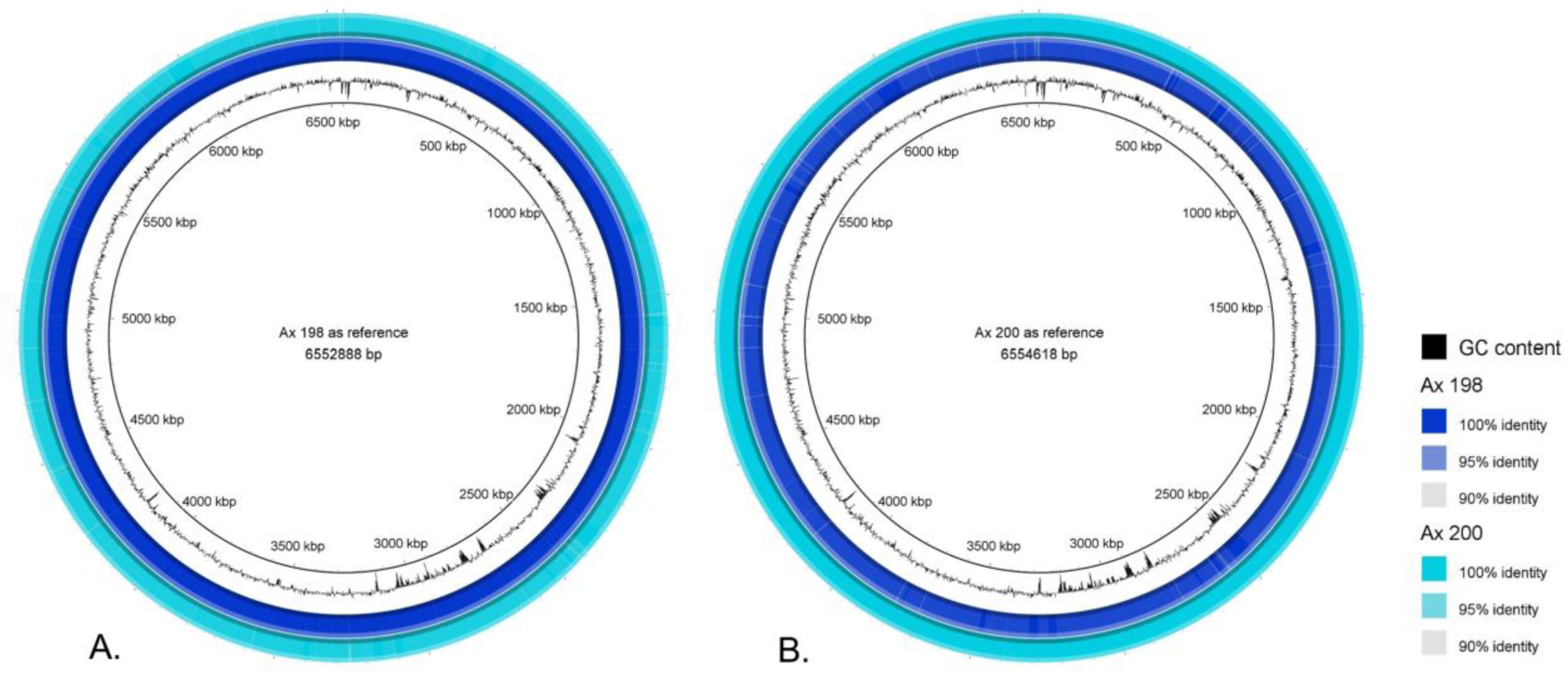
Comparative analysis of *A. xylosoxidans* Ax 198 and Ax 200 genome sequences using circular visualization. Circular diagrams, generated with BRIG (BLAST Ring Image Generator) version 0.95, display genome alignments with either Ax 198 (6552888 bp) as the reference (**A**) or Ax 200 (6554618 bp) as the reference (**B**). The innermost ring represents the GC content of the reference genome, the middle ring corresponds to the Ax 198 genome, and the outer ring to the Ax 200 genome. Genomic positions are marked in kilobase pairs (kbp). Sequence identity levels are color-coded: dark blue/turquoise for 100%, light blue/turquoise for 95%, and light grey for 90% identity with the reference genome. Regions of high similarity appear in darker shades, while gaps indicate divergence or absence, highlighting the genomic differences between the two strains.

**Figure S2.**
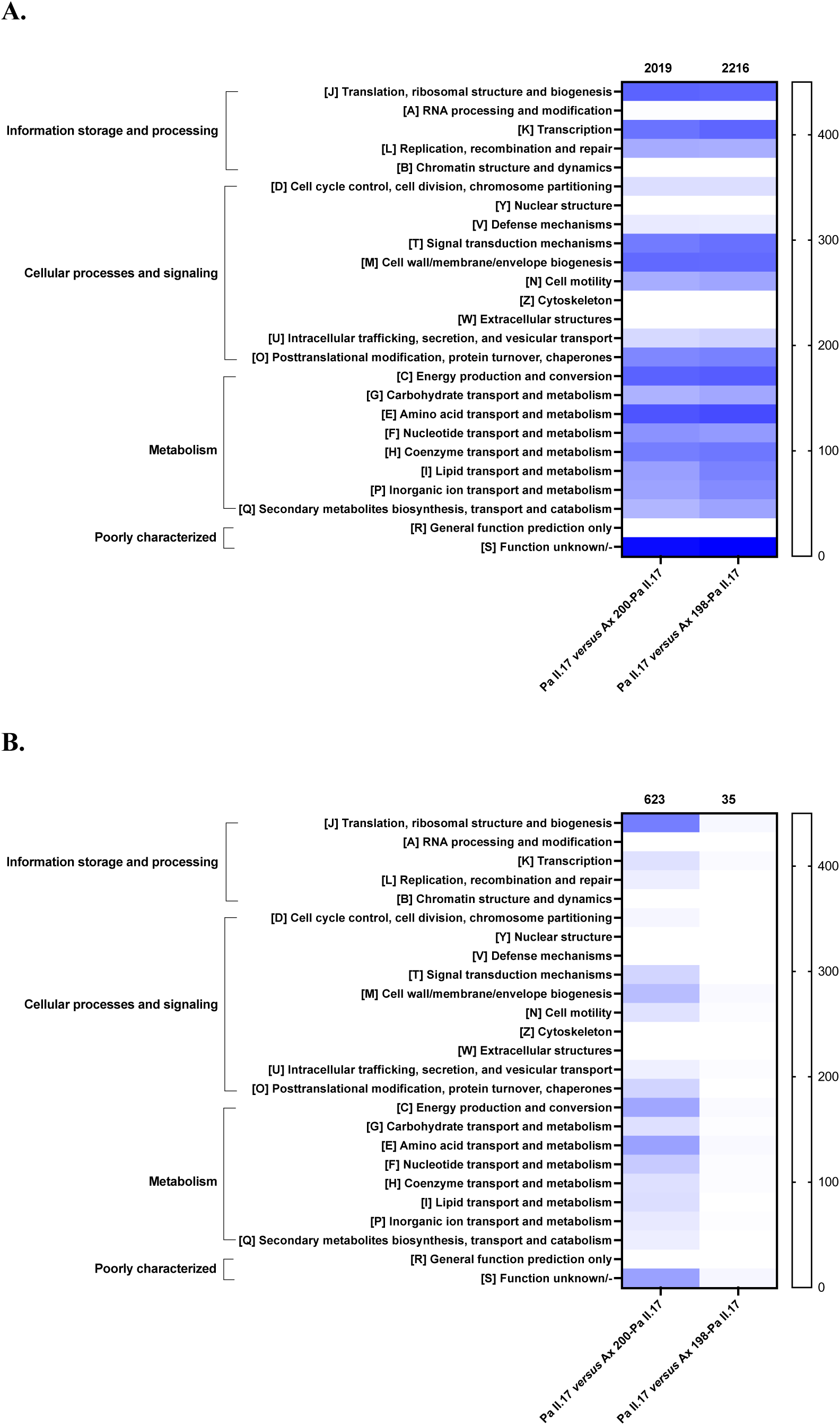
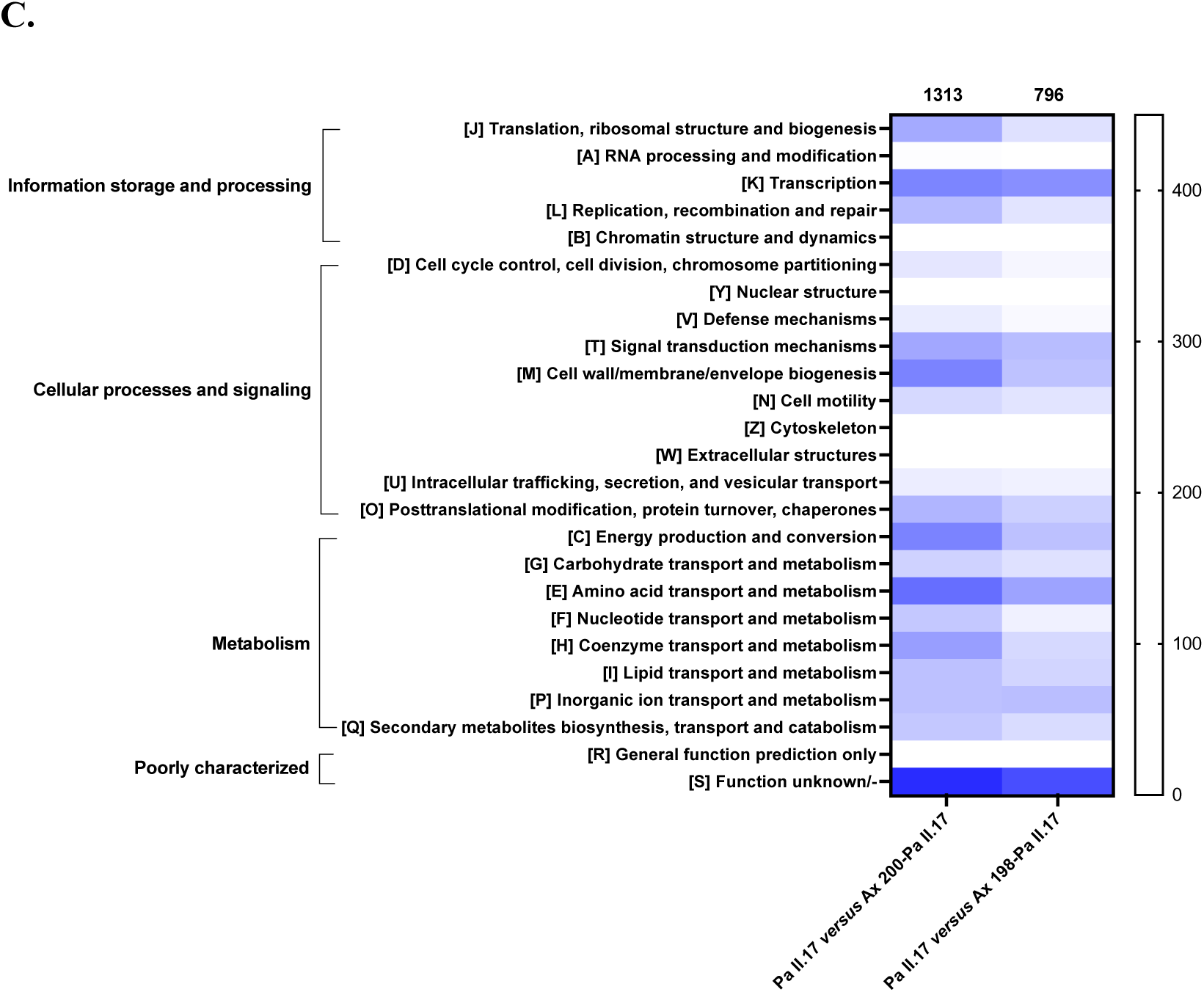
Number of *P. aeruginosa* Pa II.17 proteins with altered abundance between mono- and co-culture with *A. xylosoxidans* Ax 200 or Ax 198 according to their eggNOG database classification. **A.** Total number of proteins with altered abundance between mono- and co-culture. **B.** Proteins with significantly altered abundance between mono- and co-culture. **C.** Proteins exclusively detected in either mono- or coculture. Among the 1313 Pa proteins exclusively detected in either Pa II.17 monoculture or Ax 200-Pa II.17 co-culture, 1305 (99.4%) were unique to the Pa II.17 monoculture, while only eight were exclusive to the co-culture. Among the 796 Pa proteins exclusively detected in either Pa II.17 monoculture or Ax 198-Pa II.17 co-culture, 573 (72%) were detected exclusively in the monoculture, while 223 were unique to the co-culture.

**Figure S3.**
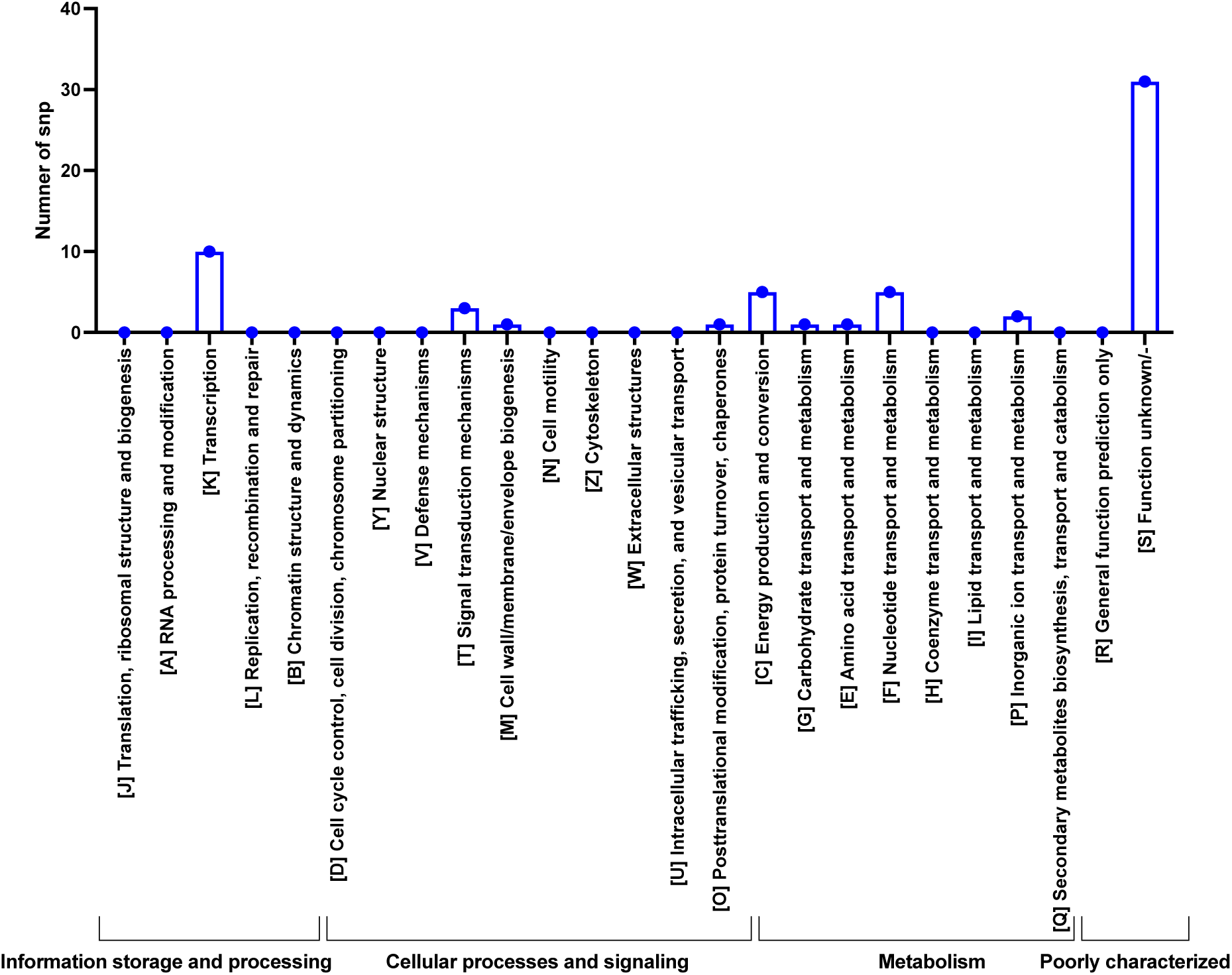
Functional distribution of SNPs between *A. xylosoxidans* Ax 200 or Ax 198 genomes. SNPs were identified using Snippy-core with default parameters, and their functional categories were annotated based on the eggNOG database.

